# Modelling Citrus Huanglongbing Spread in Scenarios involving Alternative Hosts, Vector Populations and Removal of Symptomatic Plants

**DOI:** 10.1101/2020.11.12.379743

**Authors:** Sônia Ternes, Raphael G. d’A. Vilamiu, Alécio S. Moreira, Marcelo Rossi, Tâmara T. de C. Santos, Francisco F. Laranjeira

**Affiliations:** Embrapa Informática Agropecuária, Campinas, São Paulo, Brazil; CEFET-RJ, Angra dos Reis, Rio de Janeiro, Brazil; Embrapa Mandioca e Fruticultura Cruz das Almas, Bahia, Brazil; Universidade Federal do Recôncavo da Bahia, Cruz das Almas, Bahia, Brazil

**Author notes:** Corresponding author: Francisco F. Laranjeira, Embrapa Cassava & Fruits, Cruz das Almas, Bahia, Brazil –.

**Keywords:** Asian citrus psyllid, *Diaphorina citri*, *Murraya*, HLB, epidemiology, Greening, plant disease modelling

## Abstract

Huanglongbing (HLB, ex-greening) is the most devastating citrus disease around the world. We modelled HLB spread in scenarios with different populational levels of the main alternative host (*Murraya paniculata*) and *Diaphorina citri* Kuwayama, vector of HLB associated bacteria; and removal of HLB-symptomatic plants. A compartmental deterministic mathematical model was built for representing the HLB dynamics in the Reconcavo Baiano, Bahia State, Brazil. The model encompasses delays on latency and incubation disease periods and on the *D. citri* nymphal stages. The simulations indicated that the presence of alternative hosts at low proportion would not play a crucial role in HLB dynamics in situations of poor *D. citri* management, regardless of HLB-symptomatic plants eradication. Symptomatic citrus plants contribute more to increase the HLB-incidence than the alternative host in scenarios without a suitable *D. citri* management.

## 1. Introduction

Since 2004, Huanglongbing (HLB), the most destructive citrus disease in the world (Aubert, 1992; Bove, 2006; Coletta-Filho, 2004; Da Graça, 1991), had threatened Brazilian citrus producing areas (Teixeira et al., 2005). In Brazil, HLB is associated to the bacteria *Candidatus* Liberibacter americanus (CLam) and *Ca*. L. asiaticus (CLas) (Coletta-Filho, 2004; Teixeira et al., 2005). CLas predominates in the groves (S. A. Lopes et al., 2009). The disease is incurable and affects all commercial citrus varieties. The bacteria are transmitted by the Asian citrus psyllid (ACP), *Diaphorina citri* Kuwayama (Cappor et al., 1967; Yamamoto et al., 2006). ACP is hosted by more than 50 plants from the Rutaceae family (Halbert, 2005), primarily plants of *Citrus* and *Murraya* genera. Both species also host CLam and CLas (Halbert, 2005; Lopes et al., 2006; Lopes et al., 2005). The ACP acquires the bacteria by feeding on infected plants (Bove, 2006).

The orange jasmine (*Murraya paniculata*) is an ornamental plant widely used in Brazilian backyards and urban landscapes (Laranjeira, 2011), and a host for both bacteria and their vector (Grafton-Cardwell et al., 2013). In general, plants of *M. paniculata* are considered as preferred ACP-hosts and its population are higher on this species than on *Citrus* (Teck et al., 2011). In HLB occurring areas in Brazil, the presence of orange jasmine nearby citrus groves is reported as a risk factor for HLB incidence (Aithe Junior et al., 2006). Although *M. paniculata* is the major ACP host, the pathogen does not multiply as well as in citrus plants (Lopes et al., 2010; Walter et al., 2012).

In Brazil, HLB is present in the States of Sao Paulo, Parana, Mato Grosso do Sul and Minas Gerais (Bassanezi et al., 2020). Citrus producing regions in Brazilian Northeast are free from HLB, but ACP is found (Laranjeira et al., 2018). Contingency plans to hinder HLB invasions in those regions are necessary. Such plans involve early detection and removal of infected plants. Brazilian regulation on this subject also allows local control of the orange jasmine production and trade. The role of orange jasmine in a given HLB epidemic is still controversial, because *M. paniculata* is a good host for *D. citri*, but not as suitable for *Ca.* Liberibacter spp. (Ramadugu et al., 2016). Thus, actions without scientific background such as preemptive orange jasmine eradication, could lead to ineffective and high costing results. One of the most important citrus production areas in Brazilian Northeast is the Rêconcavo of Bahia.Its citrus landscape is characterized by small orchards and a low technological level when compared with the major Brazilian citrus belt. Reconcavo of Bahia also has abundance of and frequent cycles of ACP over the year (Laranjeira et al., 2018).

The role of uncertain elements in many insect vector-related pathosystems has been investigated through mathematical modelling in order to understanding the disease dynamics (Contreras-Medina et al., 2012; Jeger et al., 2004; Stansly et al., 2014). In our study, we simulated the HLB progress in scenarios like the citrus landscape from Reconcavo (Bahia State), wherein the disease is absent, but the vector and the alternative host is widely spread. Our proposal was to verify (i) if the presence and proportion of alternative host of *D. citri* modify the HLB dynamic; (ii) if the remotion of symptomatic plants modify the disease progress; and (iii) if there was a combination of the pathosystem’s parameters that change disease status from invasive to non-invasive using the typical Reconcavo landscape (1 murraya:10,000 citrus plants).

## 2. Material and methods

The simulations were performed in a scenario with 2,700,000 citrus plants, and a fluctuation on the ACP population starting at the proportion of 0.5 insects per plant (Fig. 2 to Fig. 5). This scenario is typical of the Reconcavo Baiano citrus region. In the beginning of simulations, all the plants and insects were susceptible and the appearance of 5% ACP-infective in the total ACP-population was considered. Fluctuation of ACP infective adults have a similar behavior in both systems with and without removal of HLB-symptomatic plants

In order to test our hypotheses, a deterministic compartmental mathematical model was built comprising the main host (citrus plants), a vector population and the alternative host (orange jasmine plants). Each of the model main components were arranged in compartments.

Citrus: *S*_*c*_ – number of susceptible citrus plants; *E*_*c*_ – number of exposed citrus (latency phase); I^1^_c_ - number of asymptomatic citrus plants (incubation phase); I^2^_c_ - number of symptomatic citrus plants. The compartments for *Murraya* plants (*S*_*m*_, *E*_*m*_ and *I*_*m*_) were defined likewise. However, all the infective plants were grouped in the same group (*I*_*m*_).

The compartments related to *D. citri* (N_v_) were considered in the model as *S*_*v*_ - number of non-infective *D. citri* adults; *I*^*a*^_v_ - number of infective adults that acquired the bacteria during the adult phase; *I*^*n*^_*v*_ - number of infective adults that acquired the bacteria during the nymphal stages (Fig. 1). The total population of Citrus (*N*_*c*_), *Murraya* (N_*m*_), and ACP (*N*_*v*_) are given by:

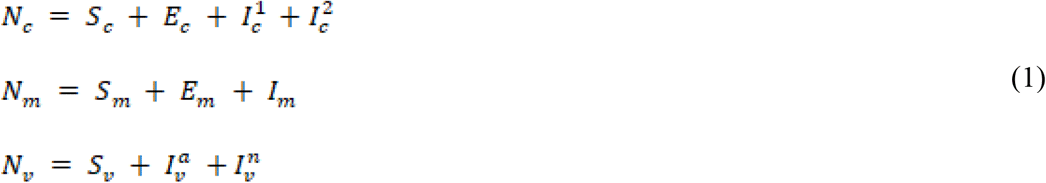

**Fig. 1.**
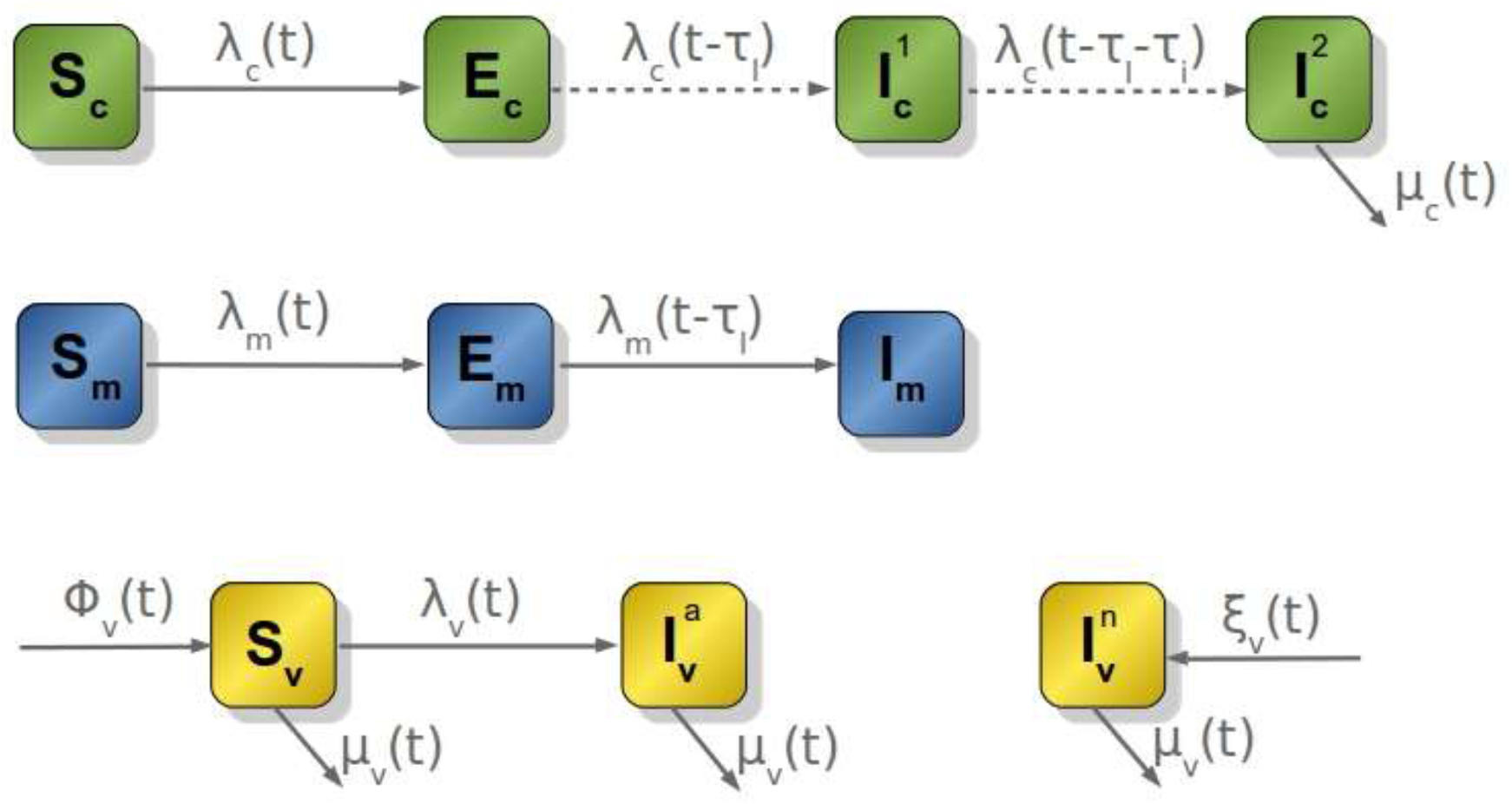
Flow diagram for the model.

Moreover, *τ*_*l*_ presented the disease latency period, *τ*_*i*_ the HLB incubation period; *τ*_*v*_ the period from egg to nymph of *D. citri*; *φ*_*v*_ the intrinsic growth function of non-infective ACP-adults; *ξ*_*v*_ the intrinsic growth function of ACP-adult that acquired the bacteria during the nymphal stages; *λ*_*c*_ and *λ*_*m*_ the force of infection from insects to citrus and *Murraya* plants, respectively, and *λ*_*v*_ the force of bacteria acquisition by ACP. The mortality rate *μ*_*(t)*_ represent discrete periodic events of infected plants removal (Fig. 1).

The model is mathematically represented by the following ordinary differential equations system:

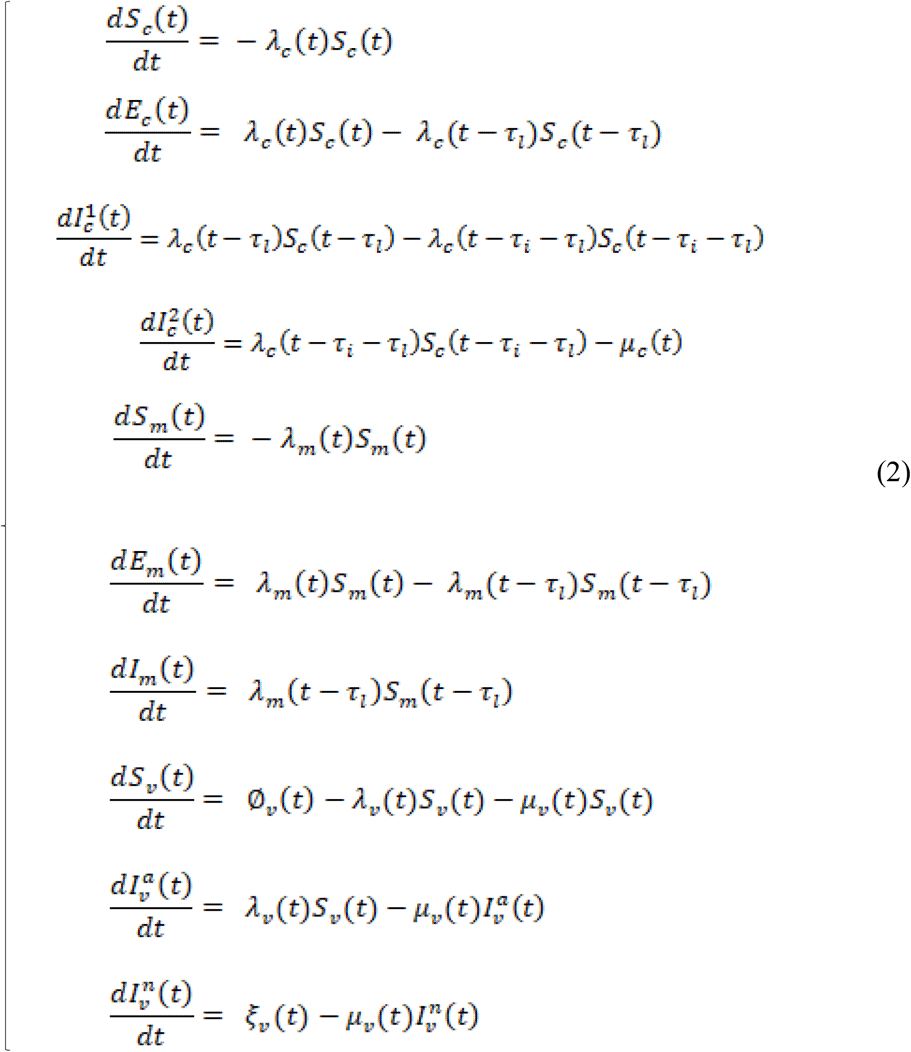

Defining *β*_*c*_ and *β*_*m*_ as the citrus and *Murraya* plants relative attractiveness, respectively (so that *β*_*c*_ + *β*_*m*_ = 1), the proportion of insects on *S*_*c*_ + *E*_*c*_, *E*_*c*_ + infective citrus, susceptible *Murraya*, and infective *Murraya* are, respectively:

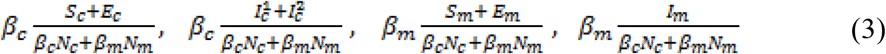

The number of new ACP-nymphs emerged from eggs layed on infected plants and reaching the adult phase will be proportional to the ACP proportion on the same infected plants, on the success rate *α* (product of the ACP reproduction rate and viability), and the total number of insects. In this way, this number can be calculated for citrus [*e*^*i*^_*c*_*(t)*] and *Murraya* [*e*^*i*^_*m*_*(t)*] plants, by

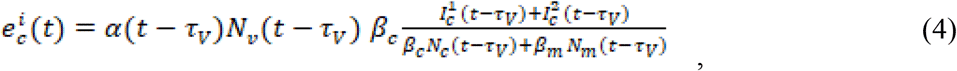

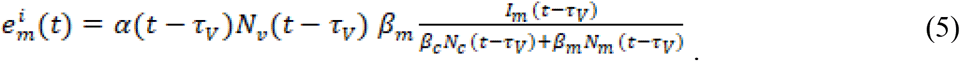

Using the definitions of *e*^*i*^_*c*_*(t)* and *e*^*i*^_*m*_*(t)* above, the intrinsic growth function *ξ* is given by:

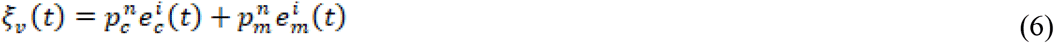

where *p*^*a*^_*c*_, *p*^*n*^_*c*_, *p*^*a*^_*m*_, *p*^*n*^_*m*_ are the acquisition probabilities (see Table 1 for parameters definitions and applied values) which are proportional to the mean bacteria concentration proportions on citrus (*C*_*c*_) and *Murraya* (*C*_*m*_):

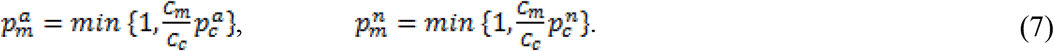

**Table 1.** Parameters for the model, their dimensions and values used in numerical simulations.

| Paramet | Definition | Value | Unit | Reference |
| --- | --- | --- | --- | --- |
| $\alpha(t)$ | Success rate (product of the reproduction rate and viability) | - | month<sup>-1</sup> | - |
| $\mu_v(t)$ | Mortality rate of adult insects | - | month<sup>-1</sup> | Equation (14) |
| $p^a$ | Probability of disease transmission from insects I<sup>a</sup><sub>v</sub> to plants | 0.035 | - | Estimate<sup>1</sup> |
| $p^n$ | Probability of disease transmission from insects I<sup>n</sup><sub>v</sub> to plants | 0.67 | - | (Nascimento, 2010) |
| $p_c^a$ | Probability of disease acquisition by I<sup>a</sup><sub>v</sub> on citrus plants | 0.43 | - | (Nascimento, 2010) |
| $p_c^n$ | Probability of disease acquisition by I<sup>n</sup><sub>v</sub> on citrus plants | 0.94 | - | (Nascimento, 2010) |
| $p_m^a$ | Probability of disease acquisition by I<sup>a</sup><sub>v</sub> on *Murraya* plants | 1 | - | Equation (7) |
| $p_m^n$ | Probability of disease acquisition by I<sup>n</sup><sub>v</sub> on *Murraya* plants | - | - | Equation (14) |
| $C_c$ | Bacteria concentration on *Murraya* plants | 3 | - | (Lopes et al., 2010) |
| $C_m$ | Bacteria concentration on citrus plants | 7.3 | - | (Lopes et al., 2010) |
| $\tau_l$ | Duration of latent phase for the disease in the plants | 1 | month | (Verbi Pereira et al., 2011) |
| $\tau_i$ | Duration of incubation phase for the disease in the plants | 6-12 | month | (Belasque et al., 2010; Bove, 2006) |
| $\tau_v$ | Duration of egg-nymph phases | 0.5 | month | (Nava et al., 2007) |
| $t_r$ | Time between successive inspections | 3 | month | (Mapa, 2008) |
| $b$ | Probing rate of hosts | 0.645 | month<sup>-1</sup> | (Laranjeira et al., 2012) |
| $\beta_c$ | Rate of psyllids on citrus plants (citrus attractiveness) | 0.05 | - | (Laranjeira et al., 2018; Santos, 2012) |
| $\beta_m$ | Rate of psyllids on *Murraya* plants (*Murraya* attractiveness) | 0.95 | - | $\beta_m = 1 - \beta_c$ |
| $\mathcal{E}$ | Human detection efficiency of symptomatic hosts | 0.47 | - | (Belasque Junior et al., 2009) |
| $c(t)$ | Psyllid mean number per plant on start of simulation | 0.41 | - | - |
| $N_c(0)$ | Initial number of citrus plants | 2,7 millions | - | - |
| $N_m(0)$ | Initial proportion from *Murraya* to citrus plants | 1:1,000–1:10,000 | - | - |

<sup>1</sup>There is no consensus in the literature about this value. However, several authors suggest that this probability is extremely low (Inoue et al., 2008; Xu et al., 1988).

The infection forces are

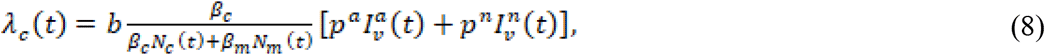

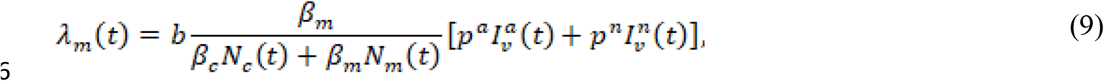

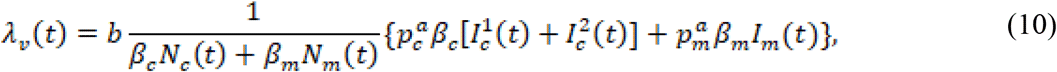

where *p*^*a*^ and *p*^*n*^ are, respectively, the probabilities of bacteria transmission from the insects (*I*^*a*^_*v*_ and *I*^*n*^_*v*_) to the plants and *b* is the probing rate of hosts. The ACP total population *N*_*v*_*(t)* was defined as a sine function with a maximum in January, based on (Yamamoto et al., 2001):

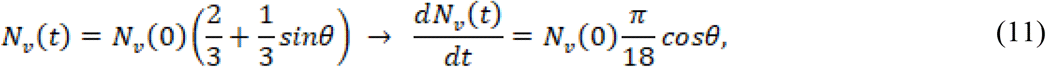

Where

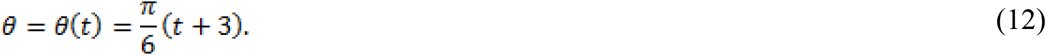

Since

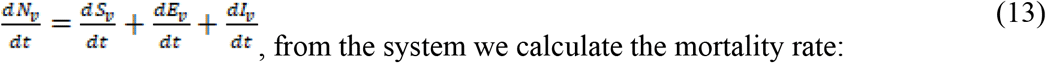

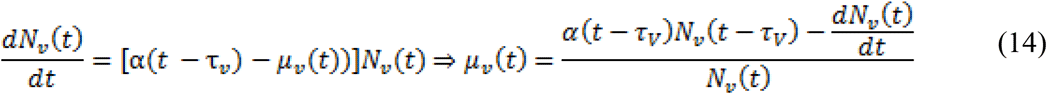

The success rate [α(t)] is fitted to get the mortality rate [*μv(t)*], which take into account the vector longevity based on (Alves, 2012), satisfying the *Nv(t)* equation [11] described above.

Since we are considering the ACP total population (*N*_*v*_) to be a scenario function, we have *S*_*v*_*(t)* = *N*_*v*_*(t)* − *I*^*a*^_v_*(t)* − *I*^*n*^_*v*_*(t)*. Furthermore, the differential equation on *S*_*v*_*(t)* can be removed from the system (1). Additionally, there is no need to define the function *φ*_*v*_*(t)* explicitly.

In Brazil, the removal of symptomatic plants is mandatory (MAPA, 2008) and the grower must periodically scouting for HLB-symptomatic plants every 3 months. Thus, the model was simulated considering periodic removal (with period *t*_*r*_) of symptomatic plants (*I*^*2*^_*c*_), when all the identified plants as symptomatic are removed based on the human detection efficiency *ε* (Belasque et al., 2010). Thus, the model determines the dynamics of compartments between two successive removals and, at each removal time, the population of symptomatic plants is proportionally reduced to *ε*.

The applied parameters in the model, their dimensions and used values in the numerical simulations are showed in the Table 1.

The yield forecast (ton) was calculated based on the mean estimated production per plant (kg) for ‘Pera’ sweet orange (*Citrus sinensis* Osb.) variety, the most planted in the Brazilian citrus belt. We considered the productivity (89.89 kg plant^−1^) reported for the season 2018/2019 (Fundecitrus, 2018). The simulations using the compartmental model gave the number of bearing trees for each year after epidemic onset. The product of number of bearing trees and production per tree was transformed in metric tons. The impact of HLB severity on yield was taken into account by multiplying the number of infected bearing trees by a HLB yield reducing factor (Bassanezi, 2018) over the first six years after HLB detection: 0% year 1; −17% year 2; −32% year 3; −44% year 4; −54% year 5; −62% year 6. From year 6 to year 10, we adopted the factor for year 6 (62%), considering that there was no symptom remission neither a recovery in plant production.

In addition to the tested scenarios, the disease incubation period was fixed at 12 months using the baseline (citrus only). Then, HLB epidemics were simulated with vector probing rates (b) ranging from 0 to 1 and average number of vectors per plant (c) ranging from 0 to 1. The result was expressed in terms of number of symptomatic plants after 10 years.

## 3. Results

Simulated scenarios with or without *Murraya* plants at the proportion 1:10,000 showed similar results for infective adults over time, regardless the removal of HLB-symptomatic plants (Fig. 2). On the other hand, in scenarios without eradication of symptomatic plants, the number of ACP infective adults would reach peaks around 60 to 70 months after epidemic onset (Fig. 2A). In scenarios where HLB-symptomatic plants were removed, the peak of ACP infective adults at that time would occur only a high proportion (1:1000) of *Murraya* plants. The other scenarios would have a great increase in the number of ACP infective adults 10 years after epidemic onset (Fig. 2B). In both scenarios, the number of ACP adults at murraya:citrus (1:1000) was higher around five years after epidemic onset (Fig. 2).

**Fig. 2.**
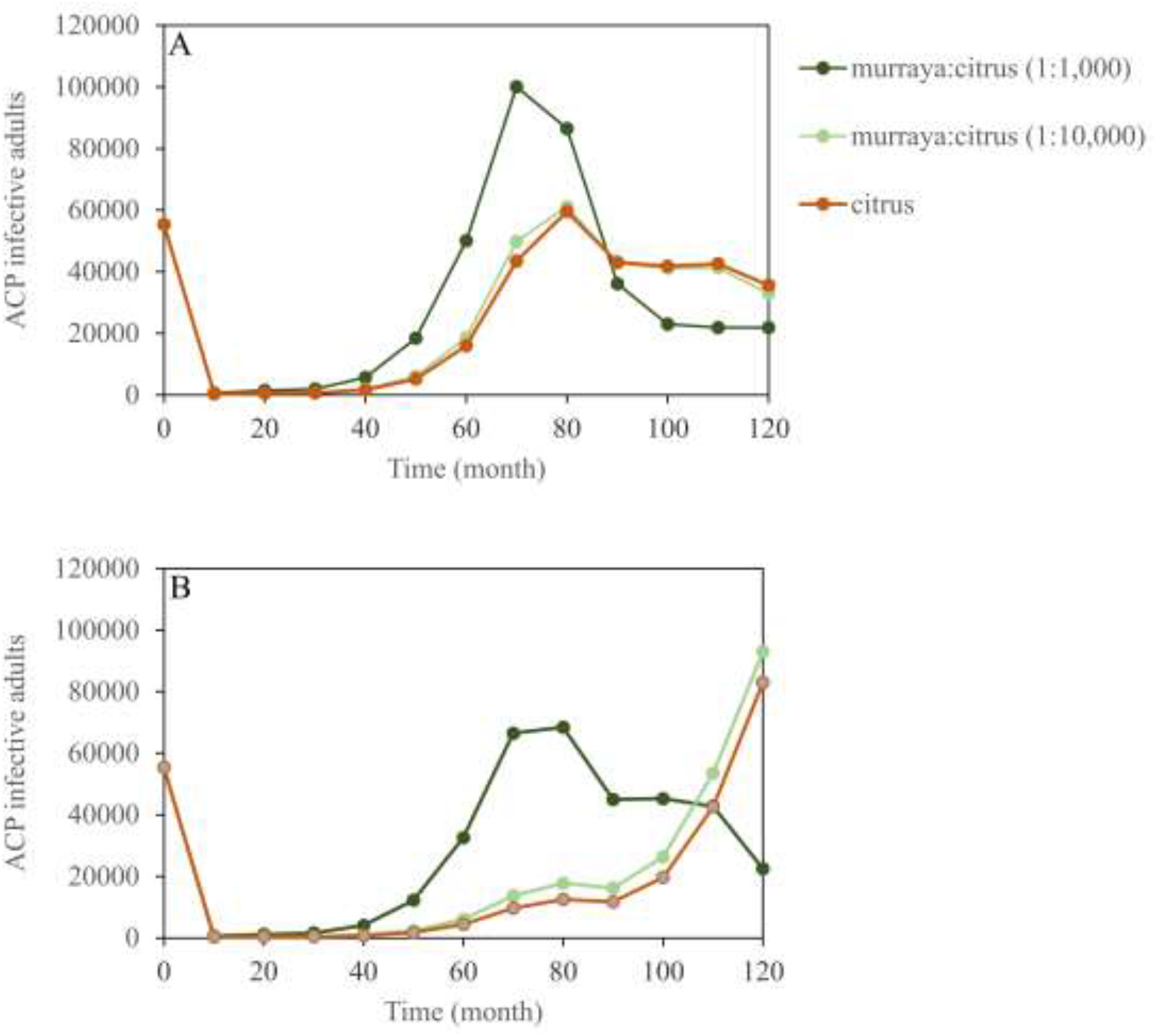
Number of ACP infective adults over 120 months of simulation in the scenarios of only citrus, murraya:citrus (1:1,000), murraya:citrus (1:10,000) for situations without (A) and with (B) removal of HLB-symptomatic plants.

The dynamics of HLB symptomatic plants did not significantly differ due the presence of murraya plants at low proportion both with or without removal of symptomatic plants. Scenarios with high number of murraya plants had more symptomatic plants over time. This difference shows up from 60 months after epidemic onset (Fig. 3). Ten years after epidemic onset, the number of symptomatic citrus plants increased significantly in simulated scenarios with murraya at high proportion and in those without murraya in scenario without removal of symptomatic plants. Landscapes with murraya:citrus (1:1,000) showed more symptomatic plants than landscapes with murraya:citrus (1:10,000) and only citrus (Fig. 3A). The progress curves of symptomatic and asymptomatic plants in the scenarios without removal were similar (Fig. 3A-B). When removal of symptomatic plants was applied, the progress curves at high proportion of murraya:citrus (1:1,000) differed from progress curves at scenarios of low proportion of murraya:citrus (1:10,000) and scenarios of only with citrus plants. (Fig. 3C-D). In the symptomatic plants’ eradication scenarios, over time there would be less symptomatic plants and more asymptomatic plants than in scenarios without removal, wherein almost 100% of the plants would be symptomatic (Fig. 3).

**Fig. 3.**
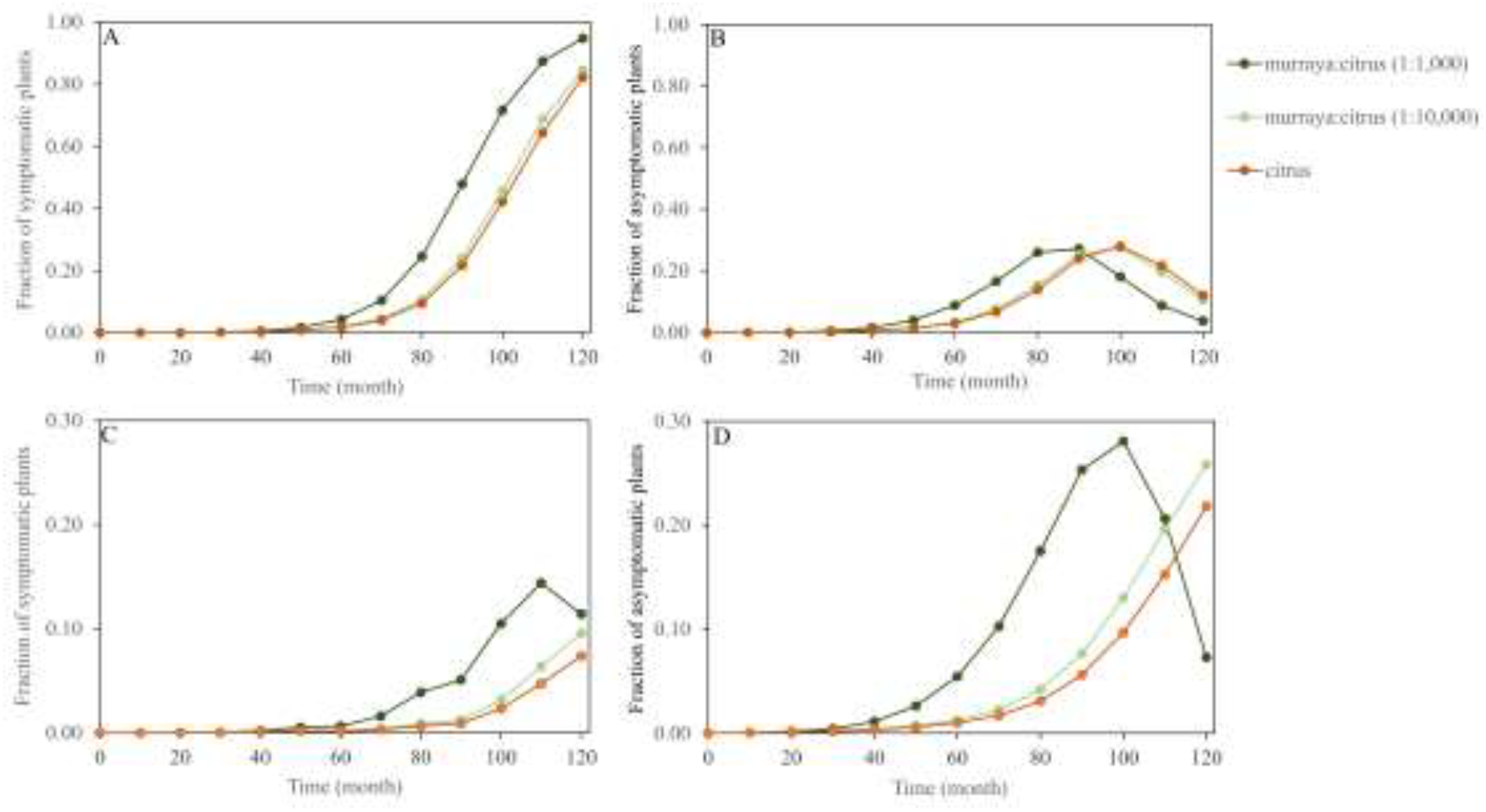
Fraction of symptomatic (A and C) and asymptomatic (B and D) plants over 120 months of simulation in the scenarios of only citrus, murraya:citrus (1:1,000), murraya:citrus (1:10,000) for situations without (A and B) and with (C and D) removal of HLB-symptomatic plants.

Regardless of eradication of symptomatic plants, the decreasing of remaining production can be noticed around 50 months after epidemic onset (Fig. 4). Without removing symptomatic plants, the progress curves would be similar. Nevertheless, four years after epidemic onset, the remaining production (%) in the proportion murraya:citrus (1:1,000) would be always less than in any other analyzed simulation. The simulations showed more than 250,000 tons 10 years after epidemic onset in all landscapes without removal of HLB-symptomatic plants (Fig. 4A).

**Fig. 4.**
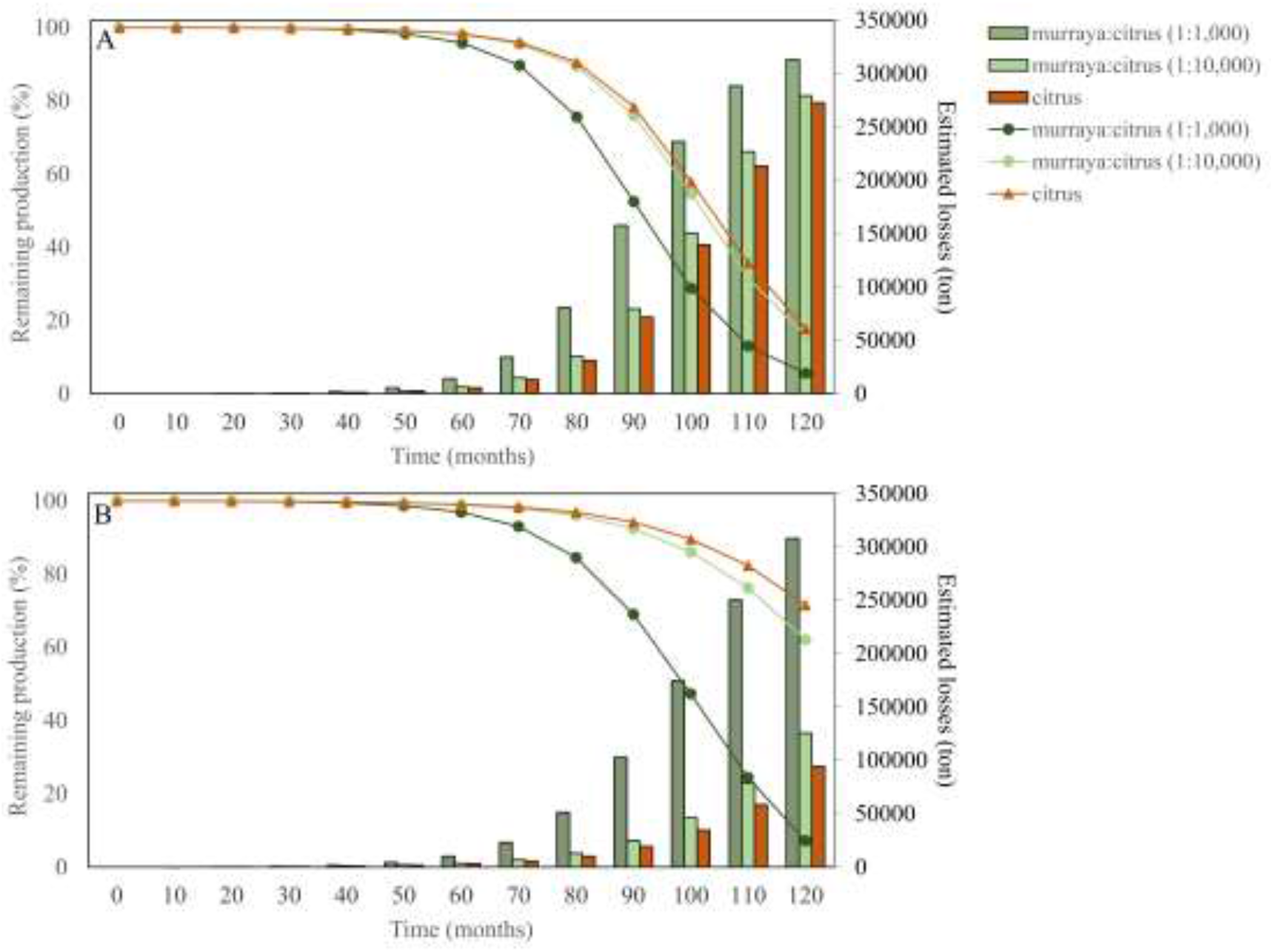
Remaining production (%) and respective estimated losses (ton) of simulation in the scenarios of only citrus, murraya:citrus (1:1,000), murraya:citrus (1:10,000) for situations without (A) and with (B) removal of HLB-symptomatic plants over 120 months since the epidemic onset date. Lines and bars dimensions are indicated by left and right y-axis, respectively.

The estimated losses for murraya:citrus (1:1,000) would reach 300,000 tons while in scenarios with only citrus or murraya:citrus (1:10,000) the estimated losses would be less than 150,000 tons (Fig. 4B).

Simulations of the combination between vector abundance fraction and vector probing rate, showed that values of 0.40 and 0.65, respectively, typical parameters for Recôncavo Baiano, and a disease incubation of 12 months, indicated that HLB epidemics in Recôncavo Baiano would be always invasive without removal the symptomatic plants and would be in a transition area with removal of symptomatic plants (Fig 5).

**Fig. 5.**
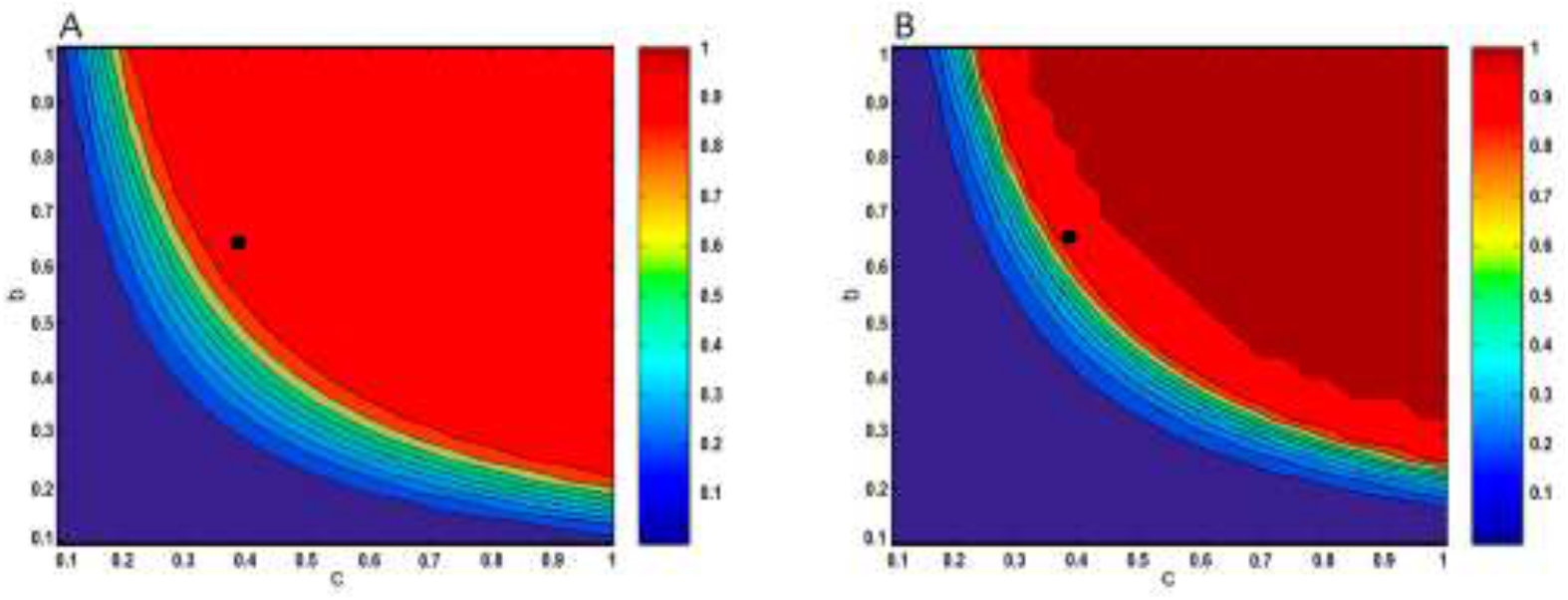
Relationship between vector abundance (c) and vector probing rate (b), and their combined effect generating invasive (red areas) or non-invasive (blue areas) epidemics. Black dots show the typical parameter combination for Reconcavo Baiano in scenario without (A) and with (B) removal considering 12 months of incubation period.

## 4. Discussion

We used a deterministic compartmental mathematical model to simulate the potential impact of an orange jasmine population on the HLB epidemic in groves of a typical smallholder citrus region in Brazil. We considered a homogeneous distribution of vectors, the latency and incubation disease periods, and the adoption of HLB management through eradication of HLB-symptomatic plants.

Our results show that HLB epidemics depend on vector abundance (mean number of psyllids per plant), the vector probe rate on the plants, the proportion of ACP alternative hosts in the susceptible plant population, and the removal the HLB-symptomatic plants. Numerical simulations suggest that in the absence of HLB management and low proportion (1:10000) of alternative hosts (i.e. orange jasmine), any control direct to those plants would not have a crucial role on HLB dynamics considering the citrus landscape used in our study. On the other hand, the adoption of removal of symptomatic plants affects the disease incidence. Nevertheless, this effect is perceived only five years after epidemic onset. This delay can arguably induce citrus growers to not adopt the removal practice as has been reported in some cases in the Brazilian citrus belt (Bassanezi et al., 2020).

Our simulations also showed that increases in the HLB incidence overlap the peak on the abundance of ACP infective adults, especially in scenario with high presence of murraya plants, regardless of removal of symptomatic plants. In addition, the losses (tons) would be more relevant in this scenario added by removal of symptomatic plants, once the losses would be a reflection of potential bacteria dissemination in murraya:citrus (1:1,000). In opposite, the losses would be less in scenarios without alternative host or when its presence is at low proportion (1:10000).

Using typical parameters of vector abundance (c) and vector probing rate (b) for Reconcavo Baiano (Table 1), we found results on the verge of invasiveness, mainly when the removal is applied. The number of infective ACP is high when the eradication of symptomatic plants is the only applied control method. Because of that, a combination of removal of symptomatic plants, reduction of vector population and reduction of urban vector refuges should stabilize HLB epidemics as long as these measures are jointly applied area-wide (Bassanezi et al., 2013; Craig et al., 2018). Conversely, should the region fail in adopting the preemptive measures, 100% of citrus plants would be infected around 10 years after the epidemic onset. That outcome would occur regardless the presence or not of alternative hosts in the region. Failure in adopting preventive practices for HLB management would have a strong impact on plants yield after 5-7 years after epidemic onset. Again, that outcome would occur regardless the presence of alternative hosts.

Policy makers could consider our results somehow counter-intuitive. Generally, policy makers consider as necessary the removal of both citrus and alternative hosts of the bacteria and ACP in backyards. Examples of that are some regulations in Mexico (Arriaga et al., 2010) or in some Brazilian cities (Paulo, n.d.). Our results point that such actions should not play a relevant role in dynamics of HLB epidemics unless the region has a high population of orange jasmine. Regarding this concern, our model could be used to investigate the relevance of *Murraya* populations and driving local decisions on the adequate policies. The disease incidence in scenarios with low proportion of orange jasmine and HLB-management by removing symptomatic plants is not different from scenarios with only citrus plants and the removal. Removal of HLB and ACP hosts as a unique measure would probably not work if applied disconnected from other disease management practices. Thereby, in a more interesting approach, orange jasmine plants could be used as biotraps (Tomaseto et al., 2016) and contribute HLB management program of plant defense agencies, as the results can be subsidize the development of pest monitoring programs in landscapes with citrus and murraya plants presence (Laranjeira et al., 2020).

## 5. Conclusions

The presence of *D. citri* alternative hosts at low proportion (i.e. 1 murraya plant:10,000 citrus plants) in a citrus landscape would not play a crucial role in the dynamics of a HLB epidemic. That holds true even without *D. citri* management and removal of HLB-symptomatic plants. Some disease impacts would show up only 5 years after epidemic onset. Symptomatic citrus plants contribute more to increase the HLB-incidence than the alternative host in scenarios without a suitable *D. citri* management.

## 6. Acknowledgments

This study was supported in part by grants from Conselho Nacional de Desenvolvimento Científico e Tecnológico - CNPq - Brazil (PQ: 309895/2016-2; PNPD: 560461/2010-0; PIBIC: 120661/2011-0).

